# SPAG6 promotes cell migration and induces epithelial-to-mesenchymal transition in luminal breast cancer cells

**DOI:** 10.1101/2022.03.24.485597

**Authors:** Jolein Mijnes, Sarah Bringezu, Jonas Berger, Carmen Schalla, Michael Rose, Sonja von Serenyi, Ruth Knüchel-Clarke, Antonio Sechi, Edgar Dahl

## Abstract

Understanding the involvement of promoter DNA methylation changes in the development of breast cancer may be highly informative for designing more effective therapeutic treatments. We recently characterized the *Sperm Associated Antigen 6* (*SPAG6*) gene, encoding a flagellar motility protein, as a potential DNA methylation biomarker for blood-based early breast cancer detection. Here we present the first study to evaluate the functional role of SPAG6 in human breast cancer. *In silico* analysis of the HumanMethylation450 BeadChip and Illumina HiSeq data of The Cancer Genome Atlas (TCGA) was performed in both normal (n=114) and breast cancer patient tissues (n=1104) to determine *SPAG6* DNA methylation and expression. Stable SPAG6 overexpressing cancer models for *in vitro* analysis were obtained by lentivirus-mediated gene delivery in T-47D, MCF-7, MDA-MB-231 and BT-549 breast cancer cells. Subsequently stable mock and SPAG6 cell lines were compared in cellular assays. In addition, involvement of SPAG6 in EMT was analysed by qPCR and immunolabeling experiments. All major molecular subtypes of breast cancer (luminal A, luminal B, basal-type, HER2-enriched) revealed a tumor-specific increased *SPAG6* promoter hypermethylation that correlated with strong reduction in *SPAG6* mRNA expression. Interestingly, a small group of luminal breast tumors exhibited *SPAG6* mRNA overexpression compared to normal breast tissue. SPAG6 overexpression caused a significant reduction (p<0.05) in colony formation in basal MDA-MB-231 and BT-549 cells. In turn, luminal T-47D cells overexpressing SPAG6 showed a significant increase in colony formation (p=0.0004) and both T-47D-and MCF-7 cells overexpressing SPAG6 exhibited a robust increase in migration speed (p<0.0001). In SPAG6-positive T-47D cells *SNAIL, TWIST1* and *Vimentin* expression was found to be significantly upregulated, while *E-Cadherin* expression was supressed. SPAG6 overexpressing T47D cells showed a typical epithelial-mesenchymal transition (EMT). This was accompanied by a nearly complete displacement of both actin and E-cadherin from cell-cell junctions. Our *in vitro* analyses give functional evidence that SPAG6 has a profound effect on colony formation, migration and intercellular junction composition in breast cancer cells. Our study is the first to show opposing SPAG6 effects in a single tumour entity depending on the molecular subtype. We propose that SPAG6 might be a key player for inducing the EMT program in luminal-type breast cancers, driving tumour progression and metastasis.

## 1. Introduction

Breast cancer remains the most frequently diagnosed cancer and leading cause of cancer deaths amongst women worldwide. Approximately 2.1 million women are newly diagnosed with this disease every year, accounting for a quarter of all cancers in women (Bray *et al*, 2018). Breast cancer is a very heterogeneous disease with diverse histopathological, genetic and epigenetic characteristics, being associated with different clinical outcomes (de Almeida *et al*, 2019; Kumar *et al*, 2013). Recent studies have shown that *de novo* promoter hypermethylation is both an early and frequent event in breast cancer development and that it is also involved in tumour progression (Brooks *et al*, 2010; Diaz & Bardelli, 2014; Tommasi *et al*, 2009; Wang & Srivastava, 2010). DNA methylation takes place at the 5-carbon position of cytosines located 5’ to guanine, the so-called CpG dinucleotide (CpG) (Dawson & Kouzarides, 2012). CpGs cluster into so-called CpG islands at promoter regions of approximately 50% of human genes (Baylin & Herman, 2000; Dawson & Kouzarides, 2012; Herman & Baylin, 2003). In breast cancer, hypermethylation of CpG islands in promoter regions is frequently observed (Baylin & Herman, 2000; Dawson & Kouzarides, 2012; Herman & Baylin, 2003) being an important mechanism for tumour suppressor gene (TSG) inactivation (Dawson & Kouzarides, 2012; Herman & Baylin, 2003; Sharma *et al*, 2010; Virani *et al*, 2012). Epigenetically inactivated TSGs are also called class II TSGs to distinguish them from TSGs that have genetic alterations and represent class I TSGs (Lee *et al*, 1991; Sager, 1997). Potential class II TSGs reported to be hypermethylated in breast cancer play important roles in e.g., cell-cycle regulation, apoptosis, DNA repair, tissue invasion and metastasis, angiogenesis and hormone signalling (Jovanovic *et al*, 2010; Tommasi *et al*., 2009). Analysing cancer-specific promoter hypermethylation of class II TSGs and understanding their contribution to breast carcinogenesis thus may give important clues towards more precise breast cancer prognosis and treatment (Jovanovic *et al*., 2010).

We previously found the gene *Sperm Associated Antigen 6* (*SPAG6*) to be hypermethylated in breast cancer and characterized it as part of the SNiPER panel for liquid biopsy-based early breast cancer detection (Mijnes *et al*, 2019). *SPAG6* is an orthologue of *Chlamydomonas Reinhardtii* Paralyzed flagella (*PF16*), which encodes for a protein localized to the central apparatus of the 9+2 axoneme involved in cilia and flagella motility (Li *et al*, 2015; Neilson *et al*, 1999; Sapiro *et al*, 2002; Teves *et al*, 2014). SPAG6 contains eight continuous armadillo repeats which are involved in protein-protein interactions (Escalier, 2006; Li *et al*., 2015; Neilson *et al*., 1999; Sapiro *et al*, 2000). Approximately half of SPAG6 deficient mice die from hydrocephalus before adulthood and males surviving to maturity are infertile due to reduced flagellar motility (Li *et al*., 2015; Teves *et al*., 2014). Interestingly, SPAG6 knockout mice revealed improper formation of the immunological synapse due to the loss of centrosome polarization and actin clearance, resulting in impaired humoral immunity (Cooley *et al*, 2016). A study on SPAG6 overexpressing COS-1 cells revealed that SPAG6 localized to structures that were positive for tubulin (Sapiro *et al*., 2000). Also, Zhang et al. found that SPAG6 decorates a subset of microtubules (Zhang *et al*, 2002), especially those surrounding the nucleus (Alciaturi *et al*, 2019).

Our original hypothesis was that SPAG6 functions, in general, as a class II TSG due to its frequent hypermethylation in breast cancer (Mijnes *et al*., 2019) and our extended analysis below). A similar situation has been described in lung cancer (NSCLC) where SPAG6 expression is also frequently lost due to promoter hypermethylation (Altenberger *et al*, 2017). However, recent data from other proliferative diseases have assigned an oncogenic role to SPAG6, e.g., in myelodysplastic syndromes (MDS) and myelogenous leukaemia (AML), where it is found to be overexpressed (Steinbach *et al*, 2006; Yang *et al*, 2015). In AML patients, *SPAG6*, amongst six other genes, could be used to sensitively monitor minimal residual disease and prognosis in AML patients (Steinbach *et al*, 2015; Steinbach *et al*., 2006). Silencing of *SPAG6* in MDS cell line SKM-1 and AML cell line K562 revealed a significant reduction in proliferation and increase in apoptosis, which was accompanied by increased expression of TP53, PTEN and various caspases, suggesting apoptosis regulation through the PI3K/Akt pathway (Yang *et al*., 2015; Yin *et al*, 2018b). In addition, liver cancer patients exhibited a significantly higher SPAG6 expression compared to normal liver tissue, which was associated with a lower 5-year survival rate (Junming, 2017). SPAG6, therefore, seems to have different roles (i.e., tumour suppressive or oncogenic functions) depending on different tissue types and the given molecular context of the cells. To date, SPAG6 expression and function has not been studied in breast cancer. We therefore aimed to investigate its role in this multifaceted and heterogeneous disease. Our data show clear evidence for a role of SPAG6 in epithelial to mesenchymal transition (EMT) and migration in luminal-type breast cancer cells.

## 2. Materials & methods

### 2.1 TCGA analysis

Infinium HumanMethylation450 BeadChip data and Illumina HiSeq data from The Cancer Genome Atlas (TCGA) breast cancer cohort were analyzed for *SPAG6*. The mean promoter methylation was calculated on basis of measured CpGs (cg12610471, cg18247055, cg05099508, cg24031355, cg10648197, cg06908778) present in the *SPAG6* gene promoter (position −348 to +165) (Shen *et al*, 2006). A Mann-Whitney U test was performed to determine the significance of differences in methylation level and gene expression between normal breast tissue and breast cancer. Spearman’s correlation analysis was used to determine correlation between *SPAG6* promoter methylation and gene expression. An overview of the clinical characteristics of the breast cancer patients is summarized in Supplementary Table 1.

### 2.2 Cell lines

Human breast cancer cell lines T-47D, MCF-7, MDA-MB-231 and BT-549 were obtained from the American Type Culture Collection (ATCC, Rockville, MD, USA), and cultured according to manufacturer’s instructions (37°C, 95% humidity, 5% CO2). The cell lines were tested for the presence of mycoplasma every month and authenticated at the start of this work (Multiplexion, Heidelberg, Germany).

### 2.3 Immunolabeling

Immunolabelling was done as already described (Gamper *et al*, 2016; Maxeiner *et al*, 2015) with minor changes. After culturing T-47D and MCF-7 cells on μ-Dishes (Ibidi, Martinsried, Germany) for 24 hours, cells were fixed using different fixation protocols depending on the antibodies or antibody combinations (Supplementary Table 2, Table 3 and Table 4). Cell nuclei was stained with DAPI. Images were collected by confocal microscopy using a LSM 700 confocal system (Zeiss) equipped with a 63x/1.3 NA objective and 350, 488, 555 and 635 nm laser lines.

### 2.4 Cloning of SPAG6-EGFP

The coding sequence for SPAG6 (transcript variant 1, NM_012443) was PCR amplified from the SPAG6-pT-Rex-DEST30 vector (Source BioScience, Nottingham, United Kingdom) and cloned into the pWPXL-EGFP expression vector (D. Trono, École Polytechnique Fédérale de Lausanne, Lausanne, Switzerland) using BamHI and MluI to generate pWPXL-SPAG6-EGFP (Supplementary Table 5). The correct sequence of SPAG6-EGFP was confirmed by DNA sequencing.

### 2.5 Production of lentivirus and cell infection

Stable overexpression was obtained by lentivirus-mediated gene delivery as previously described (Gamper *et al*., 2016; Maxeiner *et al*., 2015). Infected cells were selected by fluorescence-activated cell sorting at the Flow Cytometry Facility, a core facility of the Interdisciplinary Centre for Clinical Research (IZKF) Aachen within the Faculty of Medicine at RWTH Aachen University.

### 2.6 RNA isolation, cDNA synthesis and RT-PCR

Total RNA was isolated using the Nucleospin RNA plus kit (Macherey-Nagel, Düren, Germany), according to the manufacturer’s protocol. The RNA was eluted with 60 μl of nuclease-free water and measured spectrophotometrically with Nanodrop (Thermo Fisher Scientific). cDNA was synthesized using the reverse transcription system (Promega, Madison, USA) as described previously (Veeck *et al*, 2008). cDNAs were amplified by semi quantitative real-time PCR using the iTaq Universal SYBR green super mix (Bio-Rad Laboratories, Munich, Germany), performed in an CFX96 cycler (Bio-Rad Laboratories) according to standard procedures. All reactions were performed in triplicate. Primer sequences are listed in Supplementary Table 6. All primers spanned an exon-intron boundary. Relative mRNA expression was calculated with the comparative CT (2^-ΔΔCq^) method and normalized to WT (Livak & Schmittgen, 2001).

### 2.7 Western blot

Cell lysates and western blotting were done as previously described (Gamper *et al*., 2016; Maxeiner *et al*., 2015). Western blotting membranes were probed with different primary and secondary antibodies (Supplementary Tables 3 and 4). The signal was detected by chemiluminescence (Thermo Fisher Scientific). Equal amounts of protein (10 to 30 μg) were loaded in each lane and verified by probing with a β-actin antibody.

### 2.8 Cell proliferation and XTT assays

Cell proliferation was determined at 24, 48, 72 and 96 hours after seeding 1 x 10^5^ cells/well in 6-well plates. All measurements were done in triplicate. At the chosen time points, cells were counted with a CASY cell counter and analyser (OMNI Life Science, Bremen, Germany). For XTT assays, 1 x 10^3^ cells/well were plated in a 96-well plate. Every 24 hours, cell viability was determined using the XTT cell proliferation kit II (Roche Diagnostics, Mannheim, Germany) according to the manufacturer’s instructions. Wells containing medium without cells served as the negative controls. Measurements were done with a Tecan Infinite 2000 ELISA-reader (Tecan, Männedorf, Switzerland).

### 2.9 Apoptosis assay

For this assay, 2 x 10^4^ cells/well were plated in a 96-well plate. After 24 hours, apoptosis was quantified using the Apo-ONE homogeneous caspase-3/7 assay kit (Promega, Madison, USA) according to the manufacturer’s instructions. Cells treated with staurosporin (1 μM) served as the positive controls. Fluorescence measurements were done using a Tecan Infinite 2000 ELISA reader (Tecan, Männedorf, Switzerland).

### 2.10 Colony formation assay

1 x 10^3^ cells were plated in triplicates in a 6-well plate for up to two weeks. Colony formation was checked every day to determine the end point. Afterwards, cells were fixed and stained with 0.5% crystal violet (10% formaldehyde, 80% methanol and 10% H_2_O). The plates were then dried and imaged and the colonies counted manually.

### 2.11 Wound-healing assay

Wound healing assays were done as already described (Gamper *et al*., 2016). Briefly, 1 x 10^5^ cells were plated in silicon inserts (Ibidi, Martinsried, Germany) mounted on μ-Dishes (Ibidi) and incubated overnight at 37°C, 5% CO_2_. Cell proliferation was inhibited by treating the cells with 10 μg/ml mitomycin C for 30 minutes prior to starting the assay. Cell migration was recorded for 24 hours using a Axio observer Z1 inverted microscope equipped with a heating stage, CO_2_ controller and an Evolve EM-CCD camera driven by ZEN software (Zeiss, Jena, Germany). Average cellular speed was calculated by measuring the distance (at different locations along the wound) travelled by the wound edge before closing the wound (Gamper *et al*., 2016).

### 2.12 Statistical analysis

Statistical analyses were performed with SPSS 25.0 (SPSS, Chicago, USA) and GraphPad Prism 8.0 (GraphPad Software Inc., La Jolla, USA). Normal distribution of all data was done using the D’Agostino-Pearson normality test. Since not all data passed the normality test, statistical analyses were done using the Mann-Whitney U test. One-way ANOVA in combination with the Tukey method was used to compare three of more independent groups. p values below 0.05 were considered significant.

## 3. Results

### 3.1 Expression and promoter methylation of *SPAG6* in breast cancer molecular subtypes

We previously identified *SPAG6* hypermethylation in blood plasma of breast cancer patients suggesting a possible role of this poorly characterized gene in breast cancer biology (Mijnes *et al*., 2019). Hence, we sought to investigate the expression level and methylation status of *SPAG6* in various types of breast cancer tissues of the TCGA breast cancer cohort in more detail. In line with our previous findings, normal breast tissue showed rare *SPAG6* promoter methylation (β-value ≥0.2: 10.3%) which was significantly increased up to 95.7% in primary breast cancers (Figure 1A). In turn, *SPAG6* expression was significantly decreased in breast cancer (p<0.0001, Figure 1B). Spearman correlation analyses statistically confirmed an inverse correlation between *SPAG6* mRNA expression and DNA methylation (Spearman r: −0.41, p<0.0001). Classifying this data set by molecular subtypes, *SPAG6* hypermethylation was present in all intrinsic groups except the normal-like one (Figure 1C). Luminal A tumours showed the lowest mean methylation levels (mean β-value: 0.46) compared to luminal B (mean β-value: 0.54), HER2-enriched (mean β-value: 0.52) and basal-like tumours (mean β-value: 0.55). Interestingly, considering mRNA expression, a group of luminal A tumours (mean expression: 3.52; 75% percentile: 6.30 - maximum: 12.10) and likely also a small group of luminal B tumours overexpressed *SPAG6* compared to healthy breast tissue (mean expression: 3.64; 75% percentile: 5.23 - maximum: 6.68) as well as compared to basal-like cancers (mean expression: 2.45; 75% percentile: 2.45 – maximum: 10.26) (Figure 1D).

**Figure 1.**
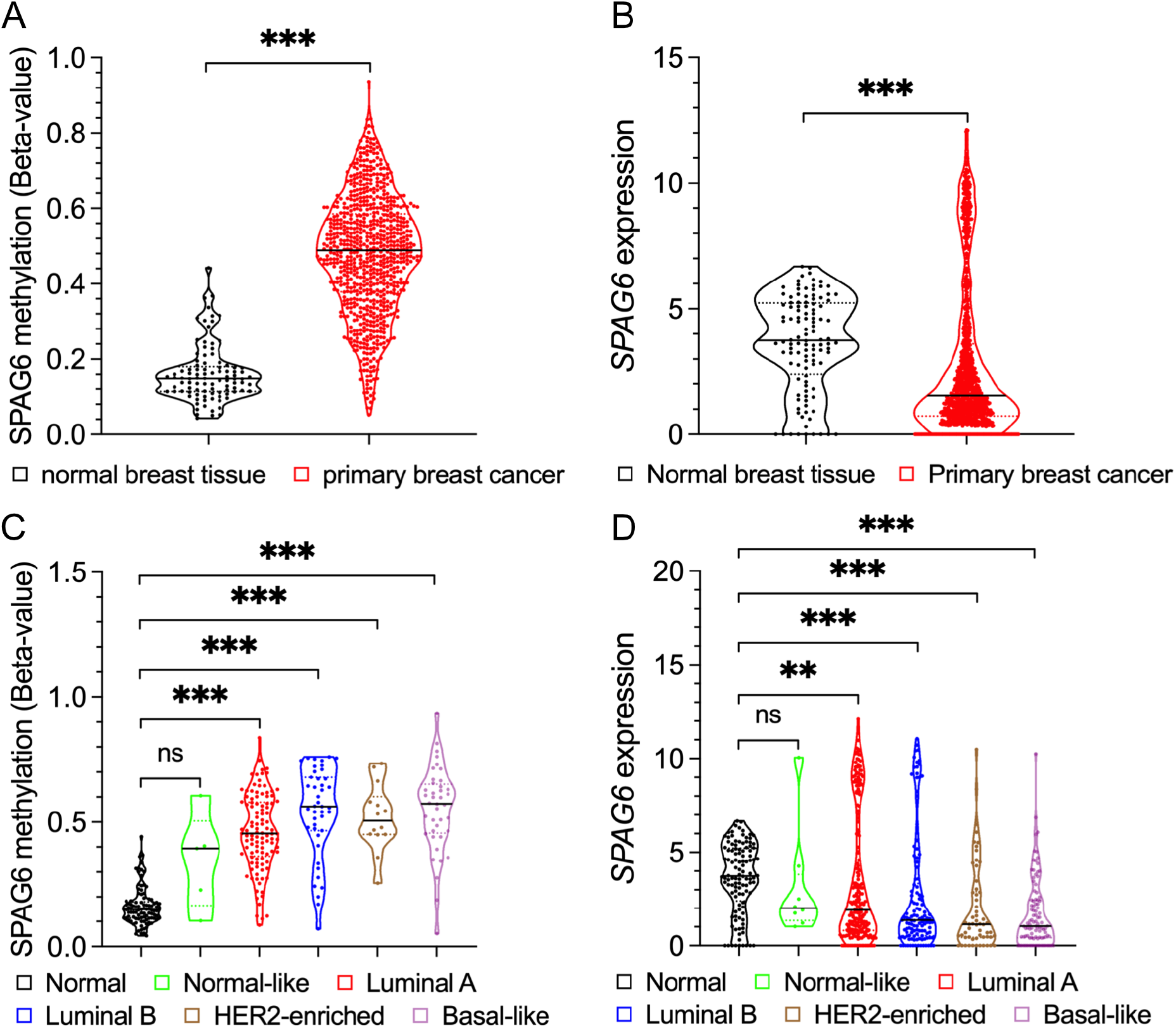
Increase of *SPAG6* promoter methylation and decrease of *SPAG6* expression in breast cancer. Methylation of the *SPAG6* gene promoter is significantly increased in primary breast cancer (A), whereas expression of *SPAG6* mRNA is significantly decreased in primary breast cancer (B). The promotor methylation frequency is also increased in the different breast cancer subtypes (C). Similarly, the downregulation of *SPAG6* mRNA expression can be observed in all molecular breast cancer subtypes (D). Notice the large spreading of the individual patients in both methylation level and gene expression. * p < 0.05, ** p < 0.01, *** p < 0.001, ns: non-significant. Grey lines indicate the median.

### 3.2 SPAG6 overexpression significantly affects colony formation capacity of luminal-type and basal-type breast cancer cells with opposite effects

Facing the divergent data of *SPAG6* mRNA expression in the TCGA breast cancer data set, we aimed to decipher the role of SPAG6 in breast cancer gain-of-function cell culture models representative for the different molecular subtypes, i.e., luminal-type (T-47D and MCF-7) and basal-type (MDA-MB-231 and BT-549) breast cancer cell lines. To this end, cells were transduced with the SPAG6-pWPXL vector, which encoded a SPAG6-GFP fusion protein, using a lentivirus-based gene-transfer system. Both mRNA and protein levels of SPAG6 were robustly expressed in all cell lines (Supplementary Figure 1).

First, colony formation capacity was measured over a period of 10-14 days. Control T-47D cells formed small but dense and well distinguishable colonies, whereas cells overexpressing SPAG6 formed much larger colonies, which were visible without the use of a microscope (Figure 2A, left panel). Quantification of mean grey scale values in crystal violet-stained colonies clearly confirmed this observation (p=0.0004, Figure 2A, right panel). During a similar period of time (14 days), however, the modulation of SPAG6 expression in MCF-7 breast cancer cells had no effect on colony formation (Figure 2B, p=0.7304). Interestingly, overexpression of SPAG6 in MDA-MB-231 cells caused a significant decrease in colony formation (Figure 2C, left panel, p=0.0061). Just like MDA-MB-231 cells, also basal-type BT549 cells overexpressing SPAG6 showed a significant reduced colony growth compared to control cells (Figure 2D, p=0.0244). It is interesting to note that SPAG6 did not exert a clear effect on cell proliferation and apoptosis, although an upregulation trend of cell proliferation could be observed especially in T47D cells (Supplementary Figure 2). Thus, our observation suggests that SPAG6 *in vitro* inhibits colony formation in basal-type cancer cells, while it supports colony formation in some luminal-type cells (i.e. T-47D cell line).

**Figure 2.**
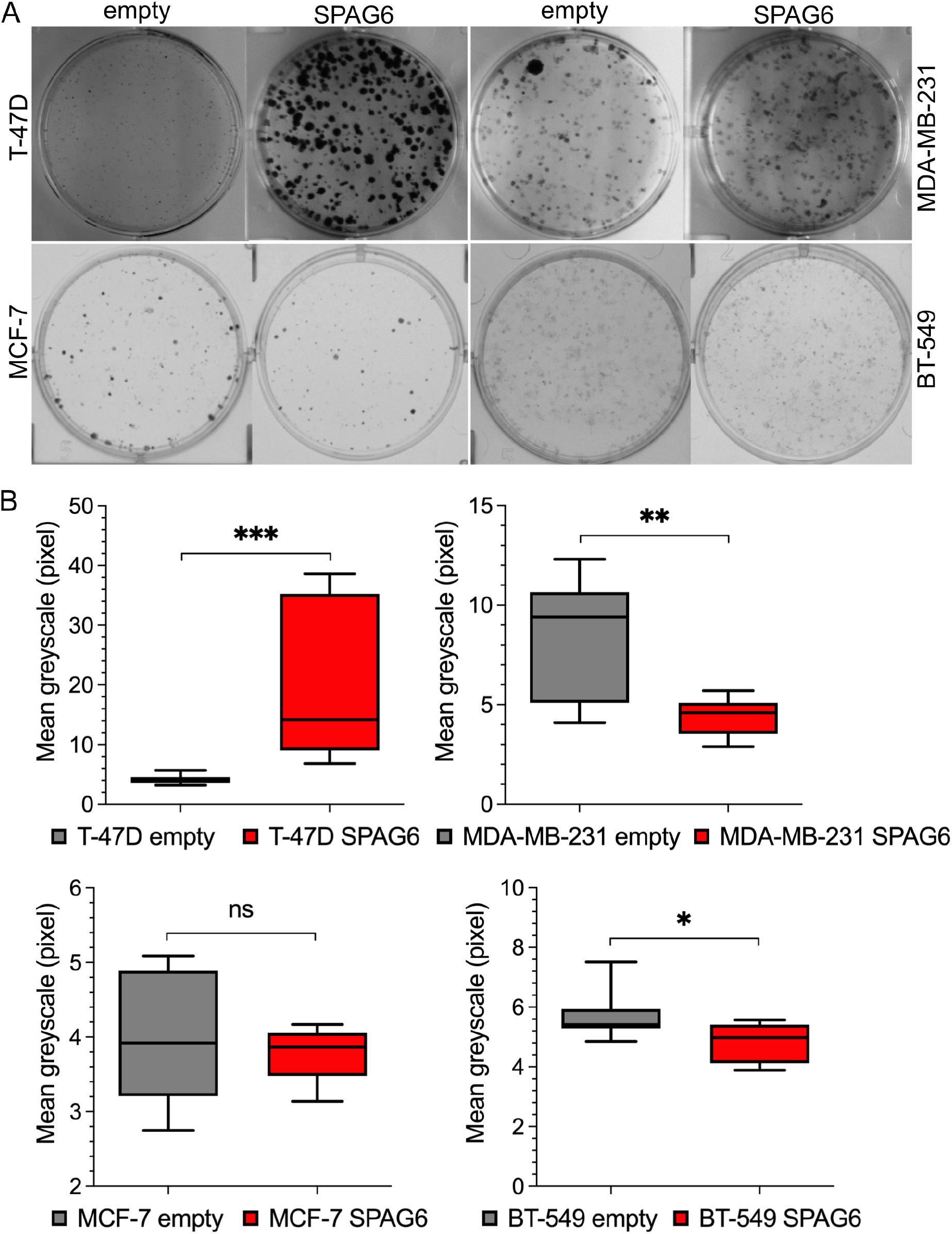
SPAG6 induces colony formation in luminal-type T-47D breast cancer cells and inhibits colony formation in basal-type breast cancer cell lines. Representative images showing colony formation in control T-47D (A), MCF-7 (B), MDA-MB-231 (C) and BT-549 (D) cells and in cells overexpressing SPAG6. Note the much bigger colonies formed by T-47D cells upon SPAG6 overexpression. Densitometry analysis of colonies after crystal violet staining is displayed in the right panels. Box plots show average grey levels. T-47D cells overexpressing SPAG6 show a significant increase in colony formation (p=0.0004), whereas MCF-7 cells overexpressing SPAG6 reveal a non-significant decrease in colony formation as compared to their control counterparts. In contrast, MDA-MB-231 and BT-549 cells overexpressing SPAG6 showed a decrease in colony formation, which was significant for both cell lines (p=0.0061 and p=0.0244, respectively). * p<0.05, ** p<0.01, *** p<0.001, ns: non-significant.

### 3.3 SPAG6 significantly promotes migration of luminal-type T-47D and MCF-7 breast cancer cells

Next, we applied classical wound healing assays to determine the effect of SPAG6 overexpression on the migration of luminal-type breast cancer cell lines. We focussed only on the luminal-type breast cancer cell lines given the larger impact of SPAG6 on their morphological features. Overexpression of SPAG6 in T-47D cells strongly increased their migration capacity observable by a much faster closing of the wound compared to wildtype cells (Figure 3A). In MCF-7 cells overexpressing SPAG6 there was no gross difference in motile behaviour compared to wildtype cells based on a visual inspection (Figure 3A). Thus, quantification of the average cell speed measured at 6 positions along each wound edge was done. Control T-47D cells migrated with a median speed of 3.9 μm/h, whereas cells overexpressing SPAG6 moved at 15.6 μm/h, thus 4x faster (p<0.0001, Figure 3B). Quantitative measurements also showed that MCF-7 cells overexpressing SPAG6 were significantly faster than their control counterparts (wildtype cells median speed: 2.0 μm/h, SPAG6 overexpressing cells median speed: 2.4 μm/h, p<0.0001, Figure 3B). Collectively, these data clearly show that SPAG6 overexpression can significantly induce cell migration capacity in luminal-type breast cancer cells.

**Figure 3.**
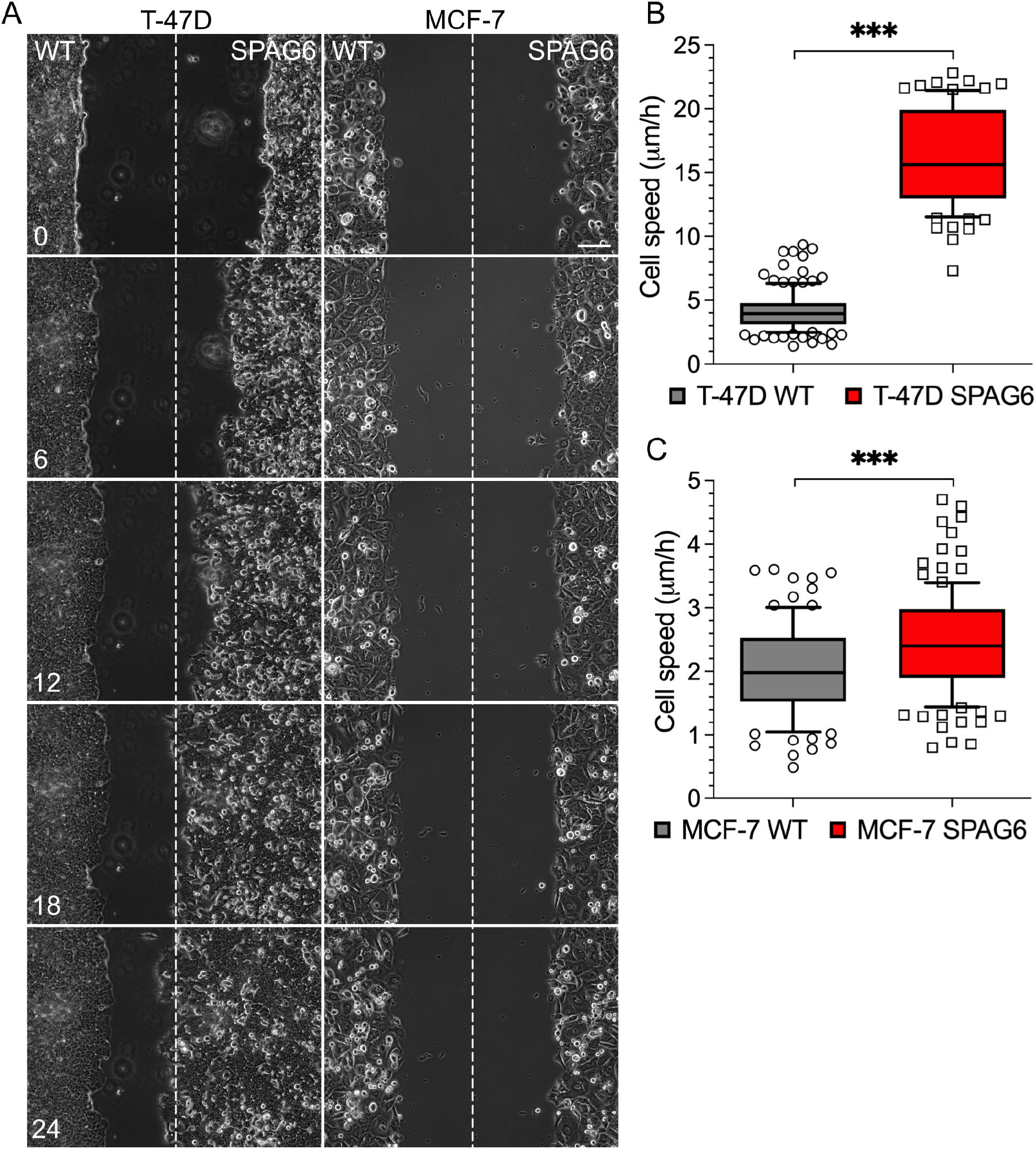
SPAG6 overexpression induces migration capacity of luminal-type T-47D and MCF-7 breast cancer cells. In (A) representative images of wound closing are shown for T-47D- and MCF-7 control and SPAG6 overexpressing cells. Dashed line indicates the middle. Cell speeds were calculated on basis of distance travelled and time (B and C). Note the more robust increase of cell migration in T-47D cells upon overexpression of SPAG6 (B). Data were generated by means of a life-imaging wound healing assay, based on two independent experiments. *** p<0.001.

### 3.4 SPAG6 overexpression induces EMT and causes a change in molecular composition of cellcell junctions in T-47D breast cancer cells

SPAG6 overexpression in T-47D cells induced prominent morphological and migration changes. For instance, T-47D cells overexpressing SPAG6 detached more easily from the cell culture vessel surface and formed several lamellipodial protrusions that were absent in wildtype cells (Figure 4A). We therefore hypothesized that SPAG6 may influence EMT in these cells. As MDA-MB-231 and BT-549 cells exhibit an intrinsic mesenchymal morphology, we proceeded with T-47D and MCF-7 cells only. The expression of well-known EMT markers such as *SNAIL, TWIST1, CDH1* (*E-Cadherin*), and *Vimentin* was determined by RT-PCR. In T-47D cells overexpressing SPAG6, the expression level of *SNAIL, TWIST1* and *Vimentin* was increased by 6.0-fold, 25.3-fold, and 2579.7-fold, respectively. In contrast, *CDH1* mRNA expression was downregulated about 10-fold compared to wildtype cells (Figure 4B). Interestingly, in MCF-7 cells overexpressing SPAG6, *SNAIL* expression was increased by 15.5-fold compared to wildtype (Figure 4C), however the expression of *TWIST1* (0.5-fold), *CDH1* (1.1-fold) and *Vimentin* (0.6-fold) remained largely unchanged (Figure 4C). The changes in *CDH1* and *vimentin* expression observed at the mRNA level could be confirmed at the protein level for both cell lines (Figure 4D).

**Figure 4.**
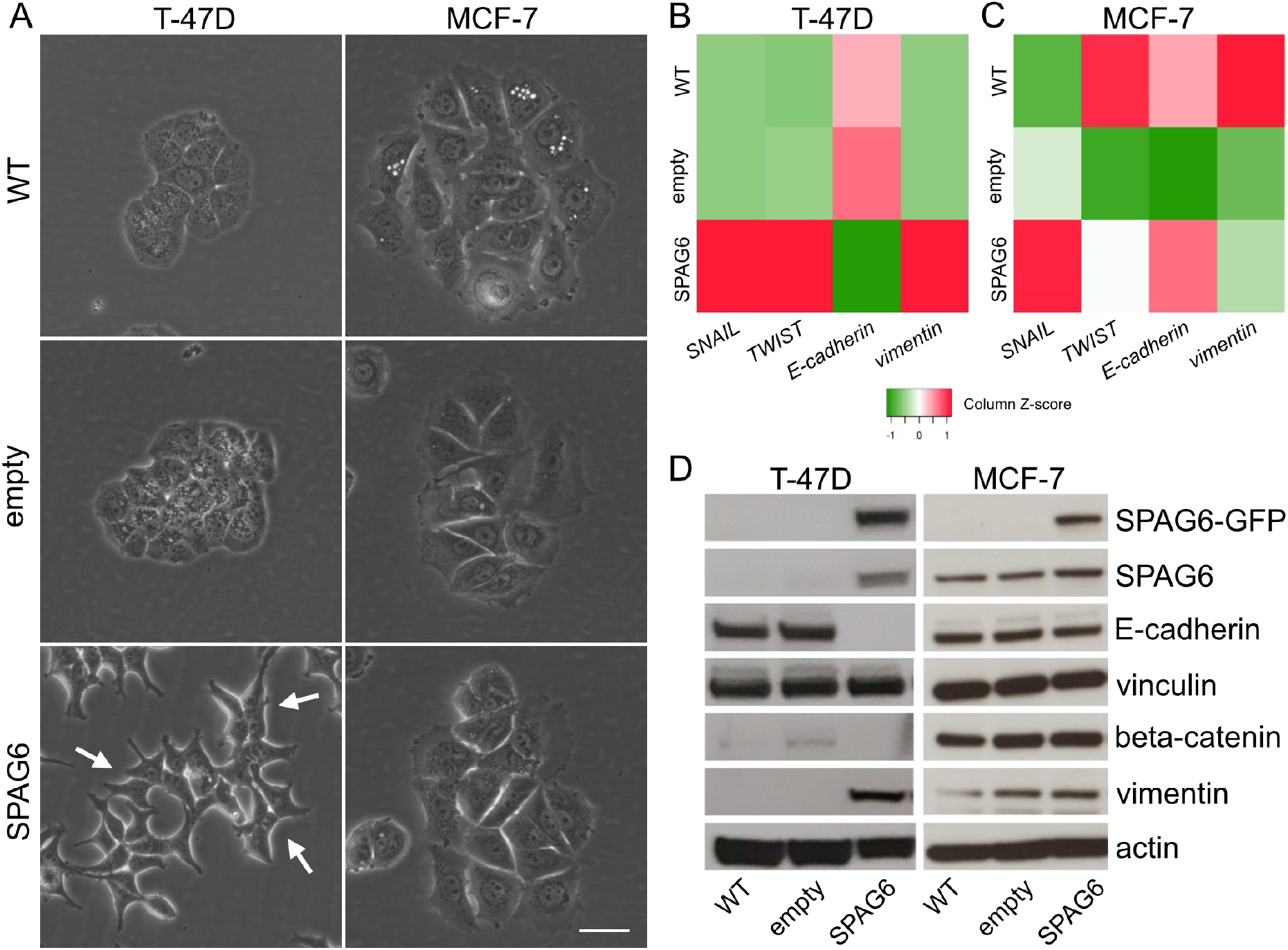
SPAG6 induces morphological changes and regulates the expression of EMT genes in T-47D breast cancer cells. T-47D cells overexpressing SPAG6 reveal a change in morphology from epithelial to mesenchymal (A, left panel, white arrows), whereas MCF-7 cells overexpressing SPAG6 do not show signs of morphological changes (A, right panel). Heatmaps showing mRNA expression levels of *SNAIL, TWIST1, E-Cadherin*, and *Vimentin* in T-47D (B) and MCF-7 (C) cells. Note the increase of *SNAIL, TWIST1, Vimentin*, and the decrease of *E-cadherin* levels in T-47D cells (B) after overexpression of SPAG6. This characteristic EMT gene signature was not induced in MCF-7 cells overexpressing SPAG6 in which *SNAIL* and *E-Cadherin* were only slightly upregulated and *Vimentin* was even slightly downregulated cells (C). Relative mRNA expression is calculated with the comparative CT (2^-ΔΔCT^) method and normalized to that of WT cells. Representative blots corroborate these finding at the protein level (D). About 20 μg of protein was loaded for each sample. Scale bar: 20 μm.

In T-47D cells, the overexpression of SPAG6 promoted the formation of less compact cell monolayers. To test this hypothesis whether SPAG6 affect cell-cell junction formation, we seeded T-47D and MCF-7 cells at high cell density and, after 24 hours, stained them with antibodies against β-catenin and E-cadherin, two specific markers of cell-cell junctions. As expected in wildtype control cells and cells expressing the empty vector, E-cadherin robustly localised at cell-cell junctions where it was found together with actin (Figure 5A). Remarkably, in T-47D cells overexpressing SPAG6, both actin and E-cadherin could hardly be found at cell-cell junctions (Figure 5A). In MCF-7 cells overexpressing SPAG6 no changes in both actin and E-cadherin localisation could be observed (Figure 5B).

**Figure 5.**
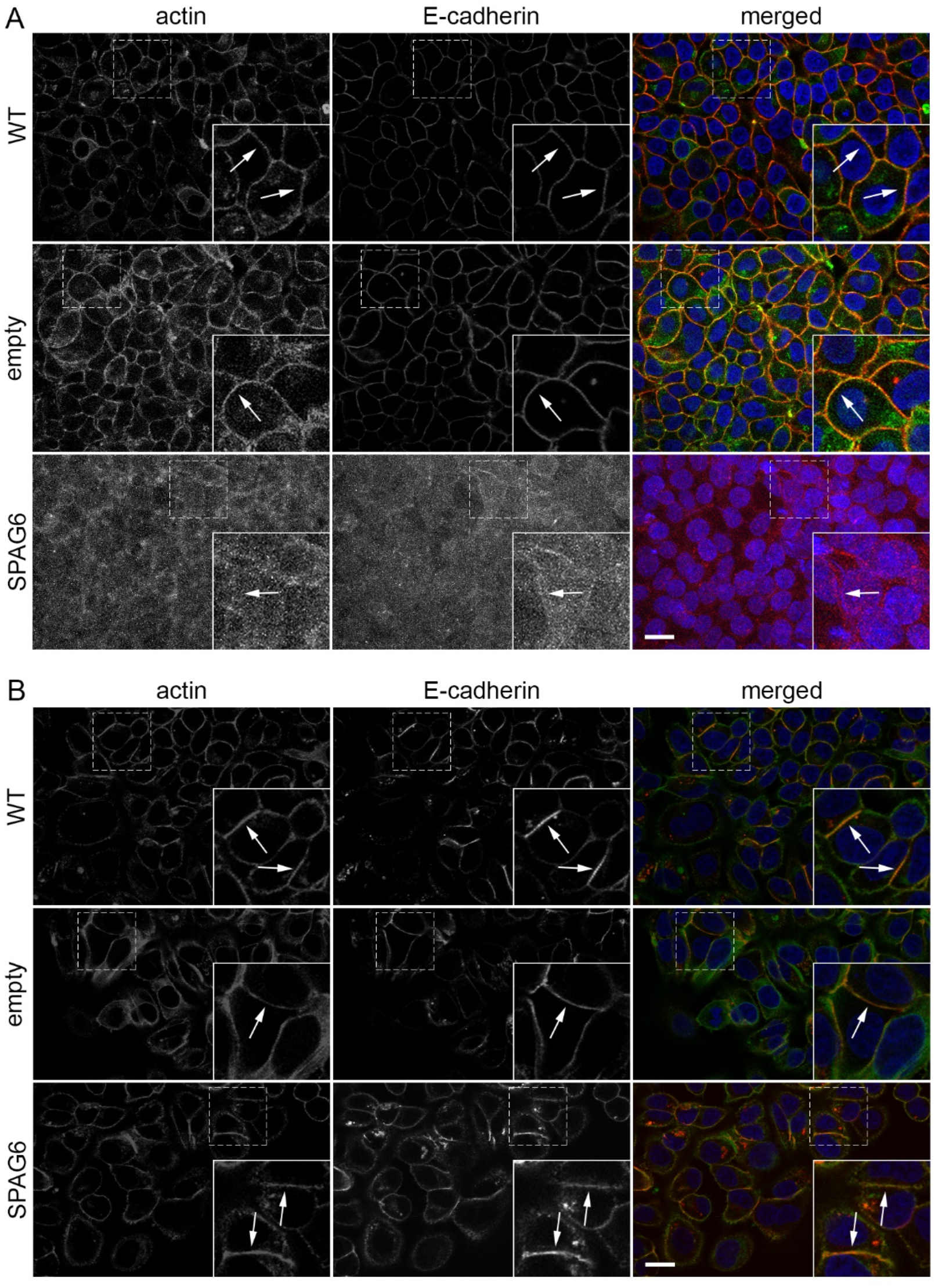
SPAG6 regulates cell-cell junction formation in T-47D breast cancer cells. Representative images showing the distribution of actin and E-cadherin at cell-cell junctions in T-47D (A) and MCF-7 cells (B). Note the co-localisation of actin and E-cadherin at cell-cell junctions in control (WT) and cell expressing an empty vector (pWPXL) (arrows in insets) in both cell types. Remarkably, the forced expression of SPAG6 causes a prominent reduction of actin and E-cadherin at cell-cell junction (arrows in insets) only in T-47D cells. Dashed boxes indicate the enlarged areas in the insets. Scale bars: 20 μm.

Since vinculin directly interacts with actin and has been found to localise at cell-cell junctions (Bays & DeMali, 2017), we determined whether the reduction of actin at cell-cell junctions in cells expressing SPAG6 also influenced vinculin localisation at these sites. Consistent with the observations above, the expression of SPAG6 in T47D, but not in MCF-7 cells, caused the displacement of vinculin and the reduction of actin from cell-cell junctions (Figure 6). Finally, given that E-cadherin is typically associated with β-catenin at cell-cell junctions (Bazzoni & Dejana, 2004; Meng & Takeichi, 2009), we also labelled T-47D and MCF-7 cells with antibodies against β-catenin and E-cadherin. Interestingly, β-catenin was found to localise at cell-cell junctions regardless of SPAG6 expression level (Figure 7A and 7B). These observations clearly indicate that SPAG6 impacts on the formation and molecular composition of cell-cell junctions and induces an EMT in T-47D breast cancer cells.

**Figure 6.**
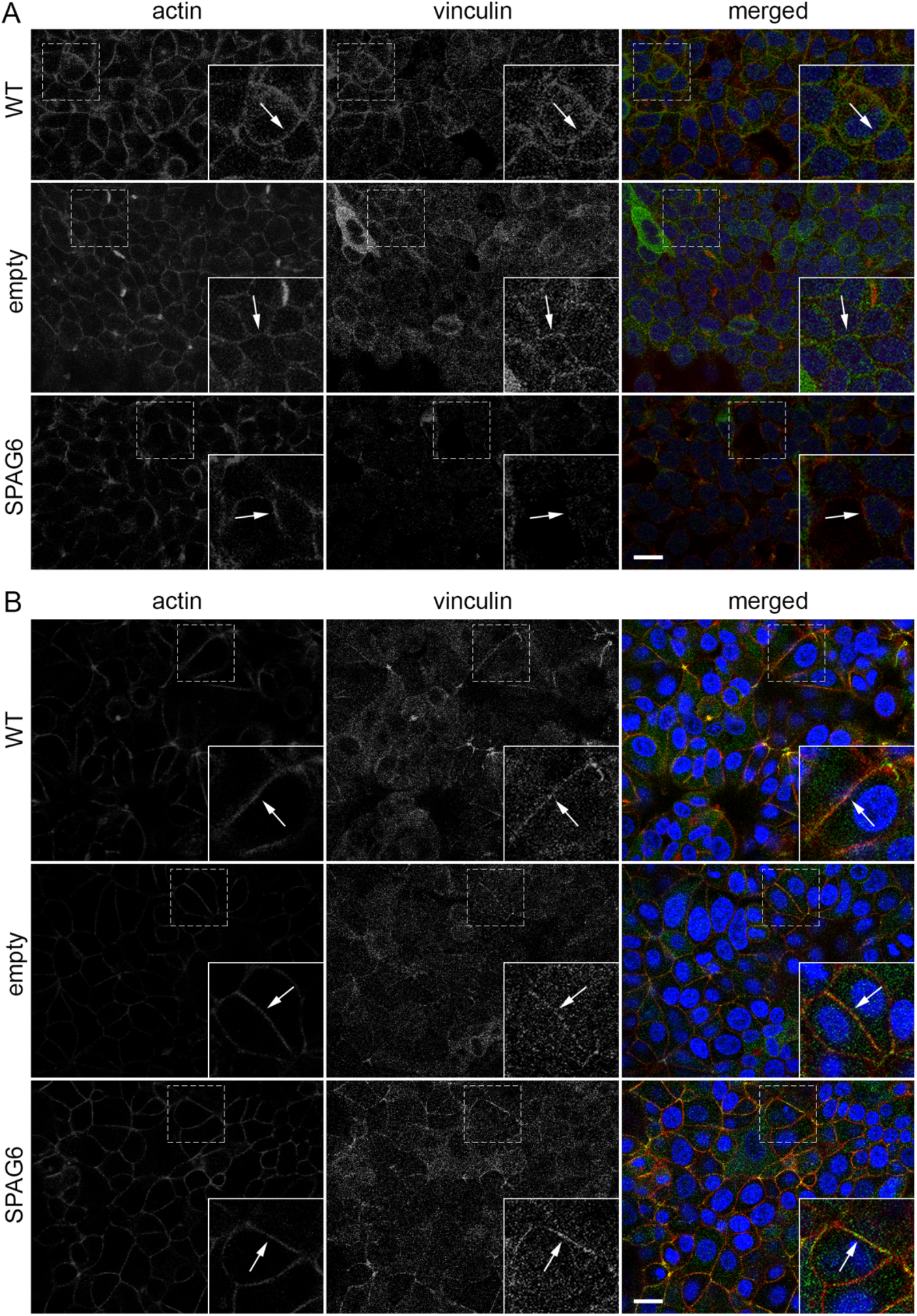
SPAG6-overexpression suppress actin and vinculin localisation at cell-cell junction in T-47D breast cancer cells. Representative images showing the distribution of actin and vinculin at cell-cell junctions in T-47D (A) and MCF-7 cells (B). Note the typical co-localisation of actin and vinculin at cell-cell junctions in control (WT) and cell expressing an empty vector (pWPXL) (arrows in insets) in both cell types. Remarkably, the increased expression of SPAG6 causes a prominent reduction of actin and vinculin at cell-cell junction (arrows in insets) only in T-47D cells. Dashed boxes indicate the enlarged areas in the insets. Scale bars: 20 μm.

**Figure 7.**
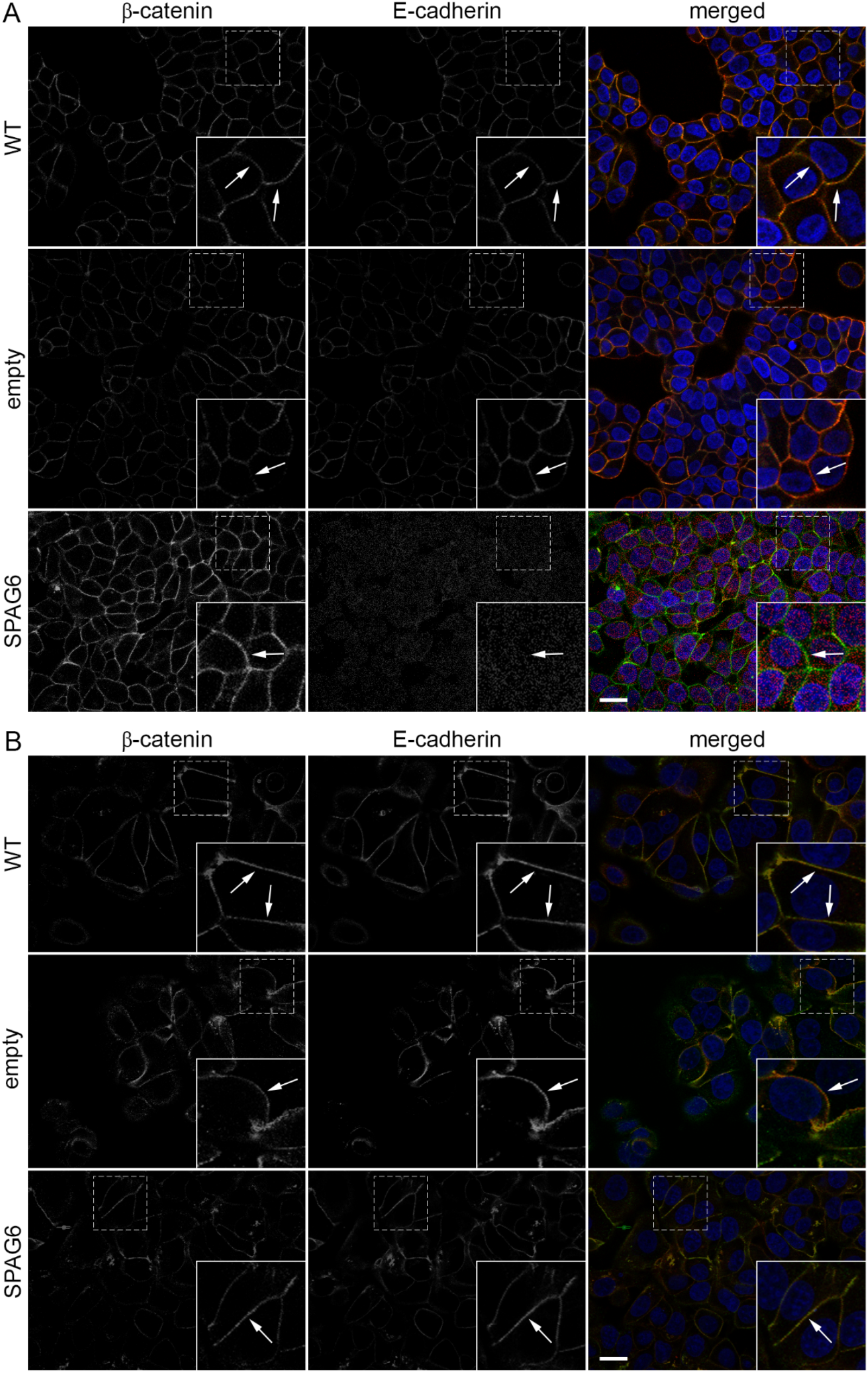
SPAG6 overexpression does not change β-catenin localization at cell-cell junctions in T-47D breast cancer cells. Representative images showing the distribution of β-catenin and E-cadherin at cell-cell junctions in T-47D (A) and MCF-7 cells (B). β-catenin and E-cadherin robustly localise at cell-cell junctions in control (WT) and cell expressing an empty vector (pWPXL) (arrows in insets) in both cancer cell lines. Notably, the increased expression of SPAG6 in T-47D cells causes a strong reduction of E-cadherin at cell-cell junctions (arrows in insets), whereas β-catenin expression and localisation are unaffected (arrows in insets). Conversely, SPAG6 overexpression does not affect β-catenin and E-cadherin localisation and expression at cell-cell junctions in MCF-7 cells (arrows in insets). Dashed boxes indicate the enlarged areas in the insets. Scale bars: 20 μm.

## 4. Discussion

Understanding the function of novel candidate genes potentially involved in the onset or progression of cancer is of crucial importance to explore new opportunities for the improvement of cancer diagnostics and therapy, particularly in the area of personalized therapies. *SPAG6* is such a candidate gene whose function in motile unicellular organisms is quite well understood, but whose function in higher organisms is still largely obscure and would not directly be associated with cancer-relevant processes. We have recently identified *SPAG6* gene as a novel potential class II tumour suppressor gene (Lee *et al*., 1991), that might be suitable for blood-based early breast cancer detection due to its abundant promoter DNA hypermethylation especially in basal-type cancer (Mijnes *et al*., 2019). The present study is the first to show that SPAG6 has a profound influence on migration and colony formation capacity of breast cancer cells. Furthermore, SPAG6 induces EMT in T-47D cells and also impacts on the formation and molecular composition of cell-cell junctions in this cell line.

In all molecular subtypes of breast cancer, except the ill-defined “normal-like” group (Bu *et al*, 2011), the *SPAG6* promoter showed a significant increase in DNA methylation levels that was associated with a significant decrease of its expression. However, a closer look at the distinct molecular breast cancer subtypes luminal A and luminal B, revealed a relatively large spreading of both methylation frequency and gene expression in these cancers. This was most pronounced in luminal A breast cancer subtype, which was clearly characterized by apparently two distinction expression patterns, i.e., a larger low expression cluster and a smaller high expression cluster (see 75^th^ percentile in Figure 1D) showing SPAG6 expression higher than that in normal breast tissue. Considering that it has been already shown for the basal-like subtype of breast cancer, that this group can be further subdivided into six molecular “sub-subtypes” including basal-like 1, basal-like 2, immunomodulatory, mesenchymal, mesenchymal stem-like and luminal androgen receptor (Lehmann *et al*, 2011; Masuda *et al*, 2013), it is possible that also the luminal A and luminal B-type breast cancer may be sub-dividable in further molecular subtypes, if we identify relevant biomarkers and SPAG6 might be such a biomarker.

Thus, our findings suggest that *SPAG6* may not have only a tumour suppressive function in breast cancer as expected based on its robust promoter DNA methylation. Based on our in vitro studies, showing that SPAG6 negatively regulated colony formation capacity of basal-type breast cancer cell lines MDA-MB-231 and BT-549, but strongly induced colony formation of luminal-type T-47D cells, it is reasonable to propose that SPAG6 might exert a dual role (oncogenic vs. tumour suppressor) depending on the molecular subtype and cellular context. This concept is also supported by the observation that cell migration, a crucial aspect of cancer development, was promoted by SPAG6 in luminal-type breast cancer cells but not in basal-type cancer. We hypothesise that a subset of luminal A (and potentially also luminal B) breast cancer cells might gain benefits for selection and survival by a SPAG6-triggered epithelial-to-mesenchymal transition process.

The notion that SPAG6 plays different roles in different cellular contexts is also supported by studies on other cancer types. For instance, *SPAG6* promoter is hypermethylated in lung cancer (NSCLC) and its expression is reduced (Altenberger *et al*., 2017), suggesting a tumour suppressive function. *SPAG6* promoter was also found to be hypermethylated in non-muscle invasive bladder cancer (NMIBC) (Kitchen *et al*, 2015; Reinert *et al*, 2011), although its methylation seemed not to be correlated with specific tumour characteristics (Kitchen *et al*., 2015). In contrast, AML patients with abundant *SPAG6* expression exhibited a poor survival (Jiang *et al*, 2019). Further studies on AML and its precursor myelodysplastic syndrome (MDS) suggested that SPAG6 regulates apoptosis through the PTEN/PI3K/AKT and TRAIL pathways (Yang *et al*., 2015; Yin *et al*, 2018a) and proliferation through the AKT/FOXO pathway (Jiang *et al*., 2019) in both diseases, suggesting that SPAG6 could be a potential therapeutic target in both AML and MDS. Liver cancer tissue also revealed higher SPAG6 expression compared to healthy tissues and patients with abundant *SPAG6* expression showed a significantly lower 5-year survival rate (Junming, 2017). In addition, *SPAG6* knockdown in the hepatocellular carcinoma cell line HCCLM3 cells caused the impairment of both proliferation and migration suggesting that SPAG6 may contribute to the development and progression of liver cancer (Junming, 2017).

Deregulation of cell motility and cytoskeleton function is a characteristic phenomenon in cancer cells contributing to their dissemination to secondary sites (Hanahan & Weinberg, 2011; Jiang *et al*, 2009; Yamaguchi & Condeelis, 2007). In tumorigenesis, cancer cells undergo molecular and cellular changes involving remodelling of cell-cell adhesions, cell-matrix adhesions and deregulation of signalling pathways leading to actin cytoskeleton remodelling (Kalluri & Weinberg, 2009; Lamouille *et al*, 2014; Yilmaz & Christofori, 2010). In this context, we clearly showed that SPAG6 has a robust impact on both T-47D and MCF-7 cells causing a significant increase of their motility. Our observations are in line with previous studies showing that SPAG6 knockdown in the hepatocellular carcinoma cell line HCCLM3 impairs cell migration (Junming, 2017), while SPAG6 knockout resulted in a similar phenotypic shift in MEFs (Li *et al*., 2015). Again, the cellular context and type of tumours must be considered when studying SPAG6 function since SPAG6 overexpression could also mediate suppressive effects as shown in neuronal cells where it impairs cell migration (Li *et al*, 2017; Yan *et al*, 2015; Zheng *et al*, 2019). Cell migration is a cyclic process which is established and maintained by signalling through Rho family GTPases, PI3Ks, integrins, microtubules and vesicular transport (Ridley *et al*, 2003). Although understanding the molecular mechanisms underlying SPAG6 function in cell migration was beyond the scope of this study, it reasonable to hypothesis that SPAG6 may be involved, either directly or indirectly, in one or more of the processes mentioned above. For instance, SPAG6 could influence the function of microfilaments, microtubules and intermediate filaments via its eight contiguous armadillo repeats (Neilson *et al*., 1999), as already shown for other proteins containing armadillo repeats including β-catenin (Fagotto, 2013; Hatzfeld, 1999; Miller *et al*, 2013; Tewari *et al*, 2010). Besides its interaction with microtubules, β-catenin is a key node in Wnt signalling and a component of actin containing junctions that links cells via cadherin proteins (Tewari *et al*., 2010).

Epithelial to mesenchymal transition is a biological process during which a cell within a tumour tissue undergoes multiple changes that enable it to acquire a mesenchymal cell phenotype, which is accompanied by changes in migratory capacity, invasiveness, and elevated resistance to apoptosis (Kalluri & Weinberg, 2009; Lamouille *et al*., 2014; Thiery *et al*, 2009). In our study, we clearly demonstrated that SPAG6 induces several EMT characteristics in T-47D cells. Along with cell elongation, formation of lamellipodia and acquisition of a front-rear polarity, all distinct features of EMT (Lamouille *et al*., 2014), SPAG6 had a great impact of cell-cell adhesions characterised by the downregulation of E-cadherin and the reduction of actin, vinculin and β-catenin localisation at these sites. Furthermore, genes known to be involved in EMT including *SNAIL, TWIST1* and *Vimentin* (Kalluri & Weinberg, 2009; Lamouille *et al*., 2014) were upregulated in T-47D cells by SPAG6. Interestingly, *SNAIL* is considered a master regulator of EMT, suppresses E-cadherin expression while increasing that of mesenchymal markers (Yilmaz & Christofori, 2010; Zeisberg & Neilson, 2009). It should be noted that MCF-7 cells did not exhibit a clear EMT phenotype although SNAIL was robustly upregulated, and their migratory capacity significantly increased by SPAG6. Since cancer cells typically exhibit different “stages” of EMT with some retaining epithelial traits while acquiring mesenchymal ones (Kalluri & Weinberg, 2009), we speculate that the different impact of SPAG6 on T-47D and MCF-7 cells produced different stages of EMT, a more complete stage in T-47D cells and a partial one in MCF-7 cells. This differential EMT behaviour could be due to the fact that MCF-7 cells already express SPAG6 and could be, therefore, less responsive to increased levels of this protein. Finally, it should be mentioned that the forced expression of SPAG6 induced collapsing of the microtubules in the breast cancer cells used in this study (data not shown). Since microtubules are important for the localisation of E-cadherin at cell-cell junctions (Dybdal-Hargreaves *et al*, 2018; Lou *et al*, 2000; Maiden *et al*, 2016; Stehbens *et al*, 2006), it is possible that SPAG6 regulate cell-cell adhesion via its impact on the function of microtubules.

Overall, we clearly show that SPAG6 overexpression differentially regulate colony formation, migration capacity and intercellular interaction of breast cancer cells in a subtype-specific manner. The impact of SPAG6 on cell migration and cell-cell junction formation suggests that this protein may play a role in inducing EMT in a defined groups of luminal breast cancer cells that has to be described at the molecular level. The clearly defined subgroup of luminal A tumours in the TCGA data set abundantly expressing SPAG6 mRNA supports this notion. In follow up studies, it will be interesting to determine in large breast tumour cohorts whether SPAG6 overexpression is associated with metastasis and unfavourable survival in this putative subgroup of luminal tumours. Future functional studies should focus on unravelling the molecular mechanisms underlying SPAG6-driven regulation of cell migration and intercellular interactions aiming at identifying novel targets for the treatment of advanced breast cancer of luminal type.

## 5. Acknowledgements

This work was supported by the Flow Cytometry Facility, a core facility of the Interdisciplinary Center for Clinical Research (IZKF) Aachen within the Faculty of Medicine at RWTH Aachen University.

## Notes

### Competing Interest Statement

The authors have declared no competing interest.

