## Supplemental Figures and Tables for "SPAG6 promotes cell migration and induces epithelial-to-mesenchymal transition in luminal breast cancer cells"

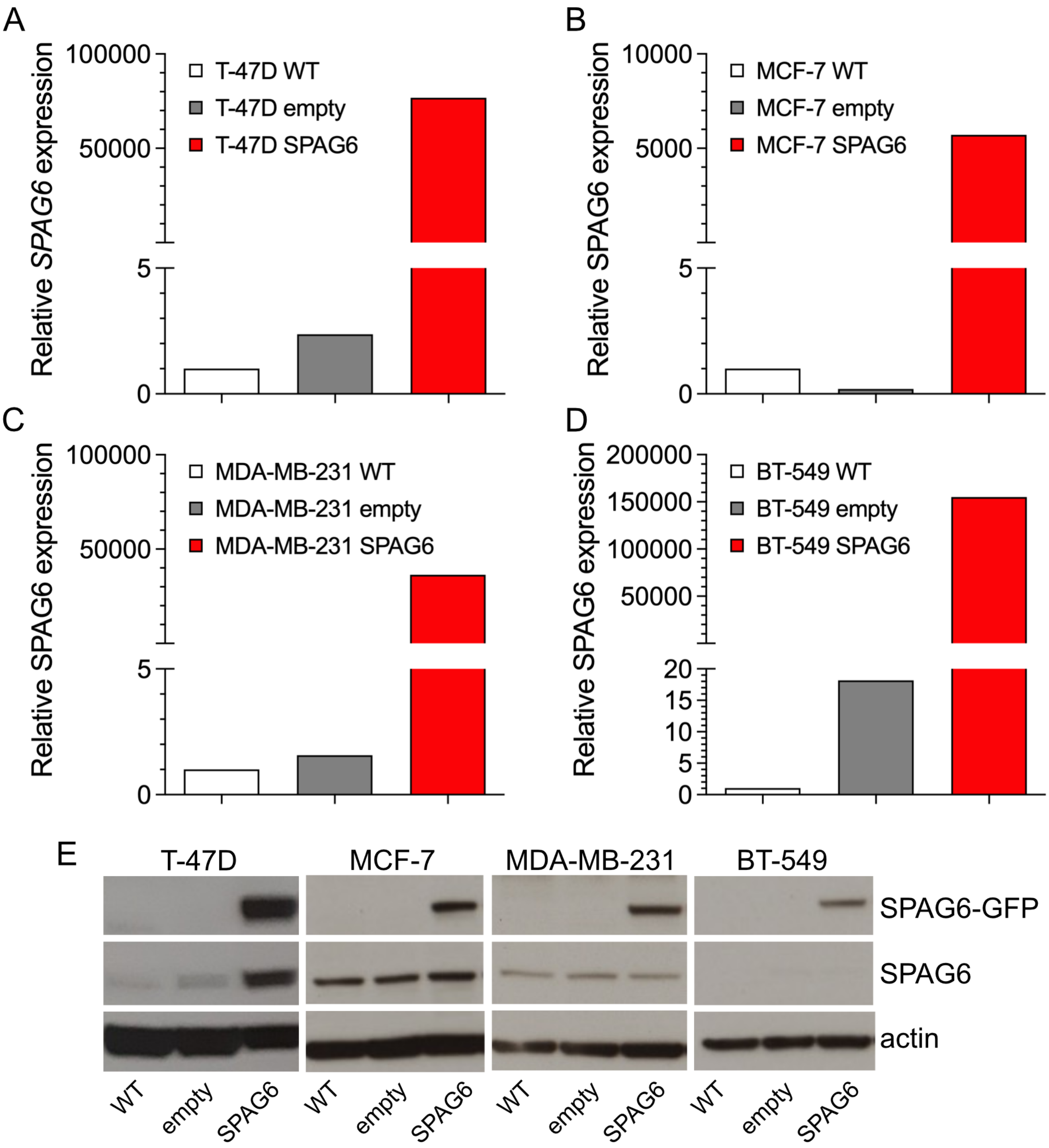


**Supplementary Figure 1. SPAG6 mRNA expression and SPAG6 protein expression is increased in all four analysed breast cell lines after lentiviral transduction.** T-47D cells show a 76758-fold increase in SPAG6 expression after transduction (A) whereas MCF-7 cells reveal a 5720-fold increase in SPAG6 expression (B). For MDA-MB-231 cells a 36460-fold increase in SPAG6 expression was observed (C) and a 155130-fold expression for BT-549 (D) as measured by qPCR. Relative mRNA expression is calculated with the comparative CT ($2^{-\Delta\Delta Cq}$) method and normalized to WT. Western blot was performed on protein lysates isolated from wildtype and transduced T-47D, MCF-7, MDA-MB-231 and BT-549 cells. Representative blots reveal strong protein bands for SPAG6-GFP at the expected size of ~75 kDa in all four analysed breast cancer cell lines (E). Some endogenous SPAG6 protein can be observed as well in T-47D, MCF-7 and MDA-MB-231 (~55 kDa). About 20 µg of protein was loaded for each sample.

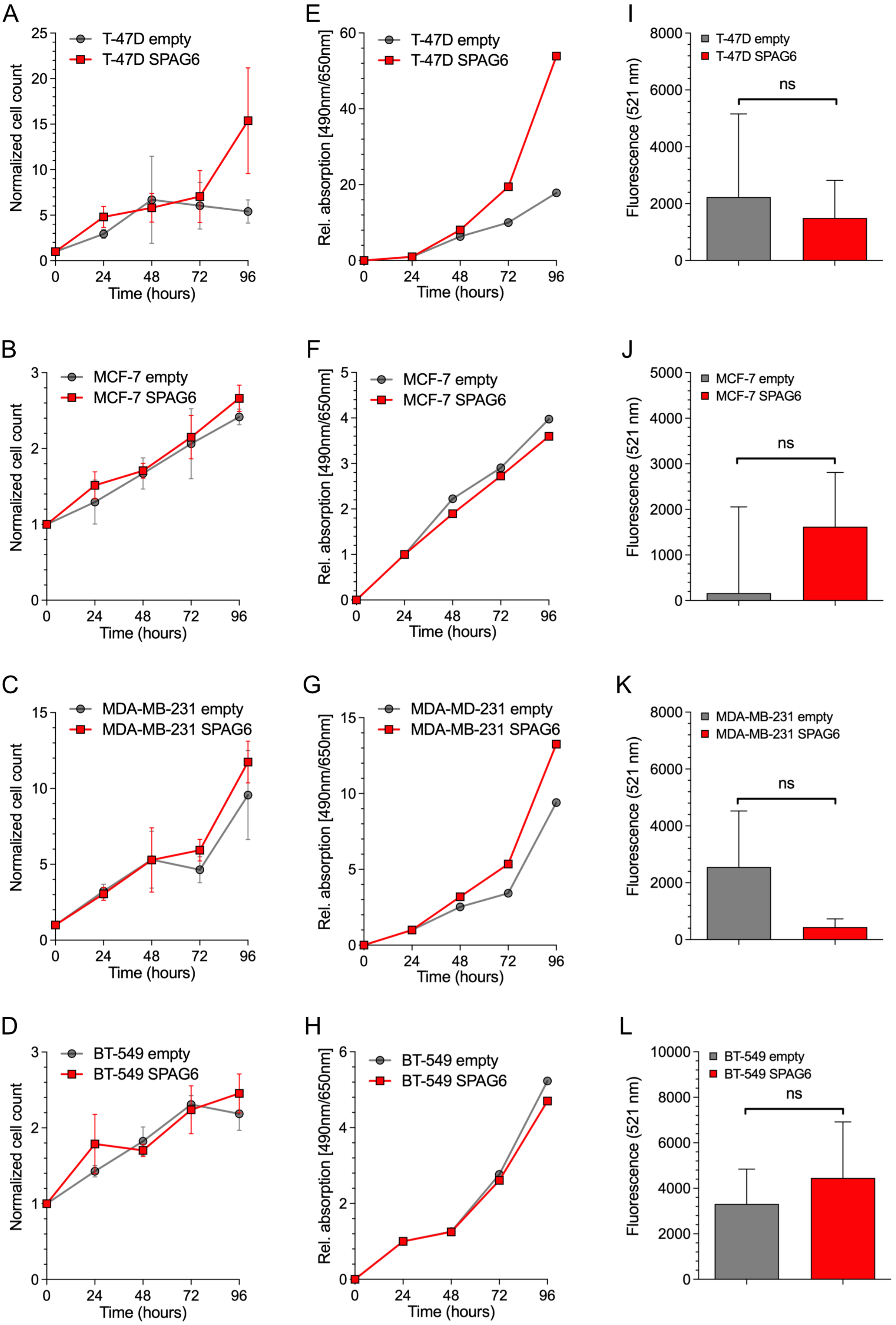
A
T-47D empty
T-47D SPAG6
Normalized cell count
Time (hours)
E
T-47D empty
T-47D SPAG6
Rel. absorption [490nm/650nm]
Time (hours)
I
T-47D empty
T-47D SPAG6
ns
Fluorescence (521 nm)
B
MCF-7 empty
MCF-7 SPAG6
Normalized cell count
Time (hours)
F
MCF-7 empty
MCF-7 SPAG6
Rel. absorption [490nm/650nm]
Time (hours)
J
MCF-7 empty
MCF-7 SPAG6
ns
Fluorescence (521 nm)
C
MDA-MB-231 empty
MDA-MB-231 SPAG6
Normalized cell count
Time (hours)
G
MDA-MD-231 empty
MDA-MB-231 SPAG6
Rel. absorption [490nm/650nm]
Time (hours)
K
MDA-MB-231 empty
MDA-MB-231 SPAG6
ns
Fluorescence (521 nm)
D
BT-549 empty
BT-549 SPAG6
Normalized cell count
Time (hours)
H
BT-549 empty
BT-549 SPAG6
Rel. absorption [490nm/650nm]
Time (hours)
L
BT-549 empty
BT-549 SPAG6
ns
Fluorescence (521 nm)

**Supplementary Figure 2. SPAG6 overexpression does not have a clear influence on proliferation and apoptosis in both luminal-type (T-47D, MCF-7) and basal-type (MDA-MB-231, BT-549) breast cancer cells.** The left panel displays data from cell counting assays (A to D), the middle panel from XTT assays (E to H) and the right panel from apoptosis assays (I to L). During the first three days, T-47D control cells and T-47D cells overexpressing SPAG6 show a similar and gradual increase in cell count. The proliferation rate of cells overexpressing SPAG6 was only somewhat higher than that of control cells during the last 24 hours of the assay (A). Similarly, control MCF-7 cells and MCF-7 overexpressing SPAG6 were characterized by a gradual increase in cell count, however no difference in cell count between the two cell lines was observed (B). Until the first 48 hours MDA-MB-231 control and MDA-MB-231 cells overexpressing SPAG6 show a similar increase in cell count, after this timepoint SPAG6 overexpressing cells show an increased proliferation compared to controls (C). Control BT-549 cells and BT-549 overexpressing SPAG6 were characterized by a gradual increase in cell count, however no difference in cell count between the two cell lines was observed (D). Triplicate measurements were performed at each time point in three independent experiments. Cell counts were normalized to cell count at plating (10.000). T-47D cells overexpressing SPAG6 show a higher metabolic activity between 48 and 96 hours after plating than control cells (E). In both MCF-7 cell type the metabolic activity was similar at any time point (F). MDA-MB-231 cells overexpressing SPAG6 revealed a higher metabolic activity between 24 and 96 hours after plating than control cells (G). In BT-549 the metabolic activity was similar at any time point (H). Multiple technical replicates (six) were performed every 24 hours for the duration of the experiment. Three independent experiments were performed. Measurements were normalized to the values of the metabolic activity measured at day 1. Overexpression of SPAG6 induced the decrease of apoptosis levels in T-47D cells (I) and MDA-MB-231 cells (K) and an increase in MCF-7 cells (J) and BT-549 cells (L). Data obtained from three independent experiments. All measurement were normalised to the basal apoptosis level. Error bars indicate one standard deviation above the mean. * $p < 0.05$, ** $p < 0.01$, *** $p < 0.001$, ns: non-significant.

### Supplementary Table I. Clinicopathological parameters of the TCGA cohort

| | N | % |
|---|---|---|
| **Age at diagnosis** | | |
| median age | 58 | |
| ≤ median | 607 | 53% |
| > median | 549 | 47% |
| Unknown | 0 | 0% |
| **Histological type** | | |
| IDC | 842 | 73% |
| ILC | 195 | 17% |
| Other | 117 | 10% |
| Unknown | 2 | 1% |
| **Tumour size** | | |
| pT1 | 291 | 25% |
| pT2 | 677 | 59% |
| pT3 | 141 | 12% |
| pT4 | 40 | 3% |
| Unknown | 7 | 1% |
| **Lymph node status** | | |
| pN0 | 535 | 46% |
| pN1 | 392 | 34% |
| pN2 | 127 | 11% |
| pN3 | 76 | 6% |
| Unknown | 26 | 2% |
| **ER** | | |
| Positive | 560 | 48% |
| Negative | 172 | 15% |
| Unknown | 424 | 37% |
| **PR** | | |
| Positive | 487 | 42% |
| Negative | 242 | 21% |
| Unknown | 427 | 37% |
| **HER2** | | |
| Positive | 103 | 9% |
| Negative | 619 | 54% |
| Unknown | 434 | 38% |

Percentages may not sum up due to rounding.

**Supplementary Table 2. Fixation and staining protocols.**

| | Fixation | | Staining | |
|---|---|---|---|---|
| | **1. step** | **2. step** | **Primary antibody*** | **Secondary antibody#** |
| **β-actin** | 1% PFA%/0,5% Tx-100 (15 min at RT) | 4% PFA (10 min at RT) | - | Phallodin-594<br>DAPI (1:1000) |
| **β-catenin** | 1% PFA%/0,5% Tx-100 (15 min at RT) | 4% PFA (10 min at RT) | anti-β-catenin (1:200) | Anti-mouse IgG-Alexa Fluor 594 (1:500)<br>DAPI (1:1000) |
| **E-cadherin** | 1% PFA%/0,5% Tx-100 (15 min at RT) | 4% PFA (10 min at RT) | anti-E-cadherin (1:200) | Anti-rabbit IgG-Alexa Fluor 647 (1:500)<br>DAPI (1:1000) |
| **Tubulin** | 4% PFA (20 min at RT) | 0,1% TX-100 in PBS (1 min at RT) | anti-YL1/2 Tubulin | Anti-rat IgG-Alexa Fluor 647 (1:500)<br>DAPI (1:1000) |
| **Vimentin** | Pre-chilled methanol (-20°C) and cells on ice | - | anti-vimentin (1:100) | Anti-rabbit IgG-Alexa Fluor 647 (1:500)<br>DAPI (1:1000) |
| **Vinculin** | 1% PFA%/0,5% Tx-100 (15 min at RT) | 4% PFA (10 min at RT) | anti-vinculin (1:200) | Anti-mouse IgG-Alexa Fluor 594 (1:500)<br>DAPI (1:1000) |

* Diluted in 1% BSA in TBS, 30-minutes incubation at RT; # Diluted in 1% BSA in TBS, 30-minutes incubation at RT in the dark; RT: room temperature

**Supplementary Table 3. List of primary antibodies.**

| Antibody | Clonality | Host | Manufacturer | Dilution |
|---|---|---|---|---|
| anti-ß-actin | monoclonal | mouse | Sigma-Aldrich, Missouri, USA | 1:2000 |
| anti-ß-catenin | monoclonal | mouse | Proteintech, Rosemont, USA | 1:200 |
| anti-E-cadherin | polyclonal | rabbit | Proteintech, Rosemont, USA | 1:200 |
| anti-tubulin | monoclonal | rat | Own production by hybridoma | undiluted |
| anti-SPAG6 | polyclonal | rabbit | Sigma-Aldrich, Missouri, USA | 1:250 |
| anti-vimentin | polyclonal | rabbit | Proteintech, Rosemont, USA | 1:100 |
| anti-vinculin | monoclonal | mouse | Sigma-Aldrich, Missouri, USA | 1:200 |

**Supplementary Table 4. List of secondary antibodies.**

| Antibody | Clonality | Host | Manufacturer | Dilution |
|---|---|---|---|---|
| anti-mouse IgG | polyclonal | goat | Dako, Glostrup, Denmark | 1:8000 |
| anti-rabbit IgG | polyclonal | goat | Dako, Glostrup, Denmark | 1:10.000 |
| anti-mouse IgG Alexa Fluor 594 | polyclonal | goat | Life technologies, Carlsbad, USA | 1:500 |
| anti-rabbit IgG Alexa Fluor 647 | polyclonal | goat | Life technologies, Carlsbad, USA | 1:500 |
| anti-rat IgG Alexa Fluor 647 | polyclonal | goat | Life technologies, Carlsbad, USA | 1:500 |

**Supplementary Table 5. Primers used for SPAG6 cloning**

| Primer name | Primer sequence (5' → 3') |
|---|---|
| SPAG6_BamHI_For | TCAGGGATCCATGAGTCAGAGGCAG |
| SPAG6_MluI_Rev | CTGAACGCGTTTGTTATTAAGTGGTTGATA |

**Supplementary Table 6. Primers used for real-time PCR and their annealing temperature (TA).**

| Primer name | | Primer sequence (5' → 3') | Cycles | TA |
|---|---|---|---|---|
| ***ß-actin*** | Forward | TGACGTGGACATCCGCAAAG | 40 | 60˚C |
| | Reverse | CTGGAAGGTGGACAGCGAGG | | |
| ***SPAG6*** | Forward | AGGGTGTACCCCAGTTGTCA | 40 | 60˚C |
| | Reverse | TTTTTACTTTTTACTTGGAGATCCTCAGAA | | |
| ***E-cadherin*** | Forward | TGCCCAGAAAATGAAAAAGG | 40 | 60˚C |
| | Reverse | GTGTATGTGGCAATGCGTTC | | |
| ***Vimentin*** | Forward | TCCACGAAGAGGAAATCCAG | 40 | 60˚C |
| | Reverse | TTCCAGGGACTCATTGGTTC | | |
| ***SNAIL*** | Forward | CTGGGTGCCCTCAAGATG | 40 | 60˚C |
| | Reverse | AGAAGGGCTTCTCGCCAGT | | |
| ***Twist*** | Forward | ACGAGCTGGACTCCAAGATG | 40 | 60˚C |
| | Reverse | CCTTCTCTGGAAACAATGACATC | | |
